# Globally stable, locally flexible: Dynamic reconfiguration of brain natural frequencies during cognitive processing

**DOI:** 10.64898/2026.05.01.721676

**Authors:** Juan J. Herrera-Morueco, Enrique Stern, Lydia Arana, Almudena Capilla

## Abstract

Neural oscillations are fundamental to brain function and cognition. Conventional analyses often rely on predefined frequency bands to assess power modulations, which may obscure finer-grained spectral variability. In this study, we focused on frequency rather than power to investigate whether the natural frequency of each brain region, typically observed at rest, represents a stable intrinsic property or dynamically reconfigures during cognitive processing. We analysed magnetoencephalography (MEG) data from the Human Connectome Project (HCP) across motor execution, working memory, and language processing tasks. Using a multivariate, data-driven spectral clustering approach, we mapped natural frequencies on a voxel-by-voxel basis without imposing predefined bands or regional boundaries. Results indicated that, while the global spatial organization of natural frequencies remained largely preserved during task engagement, specific cortical regions exhibited systematic, task-dependent shifts. In the sensorimotor cortices, the typical resting frequency of ∼24 Hz decreased to ∼6 Hz during movement preparation and at movement onset, and shifted to high-beta rhythms (∼30 Hz) following hand movement. Increased working memory demands accelerated parieto-occipital alpha/beta activity (from ∼11/16 Hz to ∼13/20 Hz) and recruited high-gamma oscillations (60 to 80 Hz) in medial temporal regions. Finally, arithmetic processing elicited a ∼5 to 15 Hz increase within the beta/gamma ranges across frontoparietal networks relative to semantic comprehension. Taken together, these findings demonstrate that natural frequencies reflect a hybrid architecture: globally stable, yet locally flexible in response to cognitive demands. Moreover, our results suggest that cognitive engagement tends to accelerate neural rhythms in functionally specialized regions, providing a more nuanced understanding of the spectral architecture of human brain function beyond conventional power- and band-based metrics.

## 1. Introduction

Neural oscillations have long been considered a fundamental mechanism of brain function, supporting communication and coordination across distributed neural populations at multiple spatial and temporal scales (Buzsáki & Draguhn, 2004; Fries, 2015; Varela et al., 2001). By temporally structuring neuronal activity, these rhythmic dynamics are thought to facilitate the integration of information across brain regions, enabling efficient large-scale network interactions underlying cognition and behavior (Singer, 2018; Wang, 2010).

Traditionally, the study of brain rhythms has relied on predefined frequency bands, with analyses often constrained to anatomically or functionally defined regions of interest (Kalamangalam et al., 2020; Keitel & Gross, 2016; Mellem et al., 2017; Niso et al., 2019). While this framework has provided valuable insights, it inherently assumes that frequency bands and cortical regions constitute homogeneous analytical units, potentially obscuring finer-grained spatial and spectral variability in neural dynamics (Buzsáki et al., 2013; Donoghue et al., 2022).

To address these limitations, a novel methodological approach has been proposed to characterize brain oscillatory activity on a frequency-by-frequency and voxel-by-voxel basis, without imposing predefined frequency bands or regional boundaries. Using this approach, Capilla et al. (2022) identified the natural frequency of each brain region—that is, the most characteristic frequency of its oscillatory activity. They further showed that the resting human brain exhibits a reproducible spatial organization of ongoing oscillatory activity, structured along two gradients of increasing frequency: one medial-to-lateral and the other posterior-to-anterior. Moreover, individual patterns of natural frequencies have been found to be remarkably robust and stable, allowing reliable identification of participants across MEG recordings acquired even four years apart (Arana et al., 2025). These findings suggest that natural frequencies capture a fundamental aspect of cortical organization that is not readily accessible through conventional band-limited spectral analyses.

However, an important open question is whether the pattern of natural frequencies, consistently observed during the resting state, is preserved when individuals engage in a cognitive task. The study of task-related modulations of oscillatory activity has largely been restricted to power-based measures. For instance, motor execution and imagery elicit robust alpha and beta power decreases over sensorimotor regions, along with low-frequency and high-gamma modulations associated with processes such as planning, execution, intention, and error processing (Bönstrup et al., 2025; Combrisson et al., 2017; Pfurtscheller & Lopes da Silva, 1999; Ramos-Murguialday & Birbaumer, 2015). Working memory tasks elicit load- and phase-specific power modulations, characterized by frontal theta increases during maintenance, posterior alpha/beta changes during encoding and updating, and high-gamma recruitment in medial temporal regions during active manipulation (Carver et al., 2019; Heinrichs-Graham & Wilson, 2015; Jensen & Tesche, 2002; Mirjalili et al., 2025; Semprini et al., 2020). Similarly, language comprehension involves delta and theta oscillations tracking syntactic and prosodic structure, while beta power suppression in frontal–temporal regions has been suggested to reflect semantic unification. In contrast, arithmetic processing additionally engages frontoparietal beta–gamma interactions (Bastiaansen & Hagoort, 2006; Dekydtspotter et al., 2025; Gross et al., 2013; Lam et al., 2016; Lewis & Bastiaansen, 2015; Lu et al., 2023; Prystauka & Lewis, 2019).

In this study, we propose to focus on frequency instead of power to investigate whether natural frequencies are dynamically reorganized during active cognitive processing. We hypothesize that the characteristic resting-state pattern of natural frequencies will be preserved, while task-specific spectral reconfigurations will emerge in specific regions related to each cognitive function. To this end, we analyzed MEG data from the Human Connectome Project (HCP) while participants completed motor execution, working memory, and language processing tasks. K-means clustering was applied to source-level power spectra to identify recurrent spectral patterns and generate individual brain maps of natural frequencies, which were subsequently compared voxel-wise across conditions to assess task-related reconfigurations. By adopting an analysis strategy that avoids predefined frequency bands and region-of-interest constraints, the present work aims to provide high spatial and spectral resolution insights into how cortical oscillatory dynamics are organized and reshaped by cognitive demands.

## 2. Material and methods

### 2.1. Participants

Data were obtained from the 1200 Subjects Data Release of the HCP (WU-Minn Consortium Human Connectome Project, 2017a), an open-access database which provides anonymized task MEG recordings and T1-weighted Magnetic Resonance Images (MRIs) (Larson-Prior et al., 2013; Van Essen et al., 2013).

The study sample consisted of 61 healthy young adults for the motor task, 83 for the working memory task, and 82 for the language processing and arithmetic task. Gender was balanced, and age was anonymized using predefined ranges. Participants’ demographic characteristics are summarized in Table 1. All procedures complied with the Declaration of Helsinki and were approved by the institutional ethics committees.

**Table 1.**
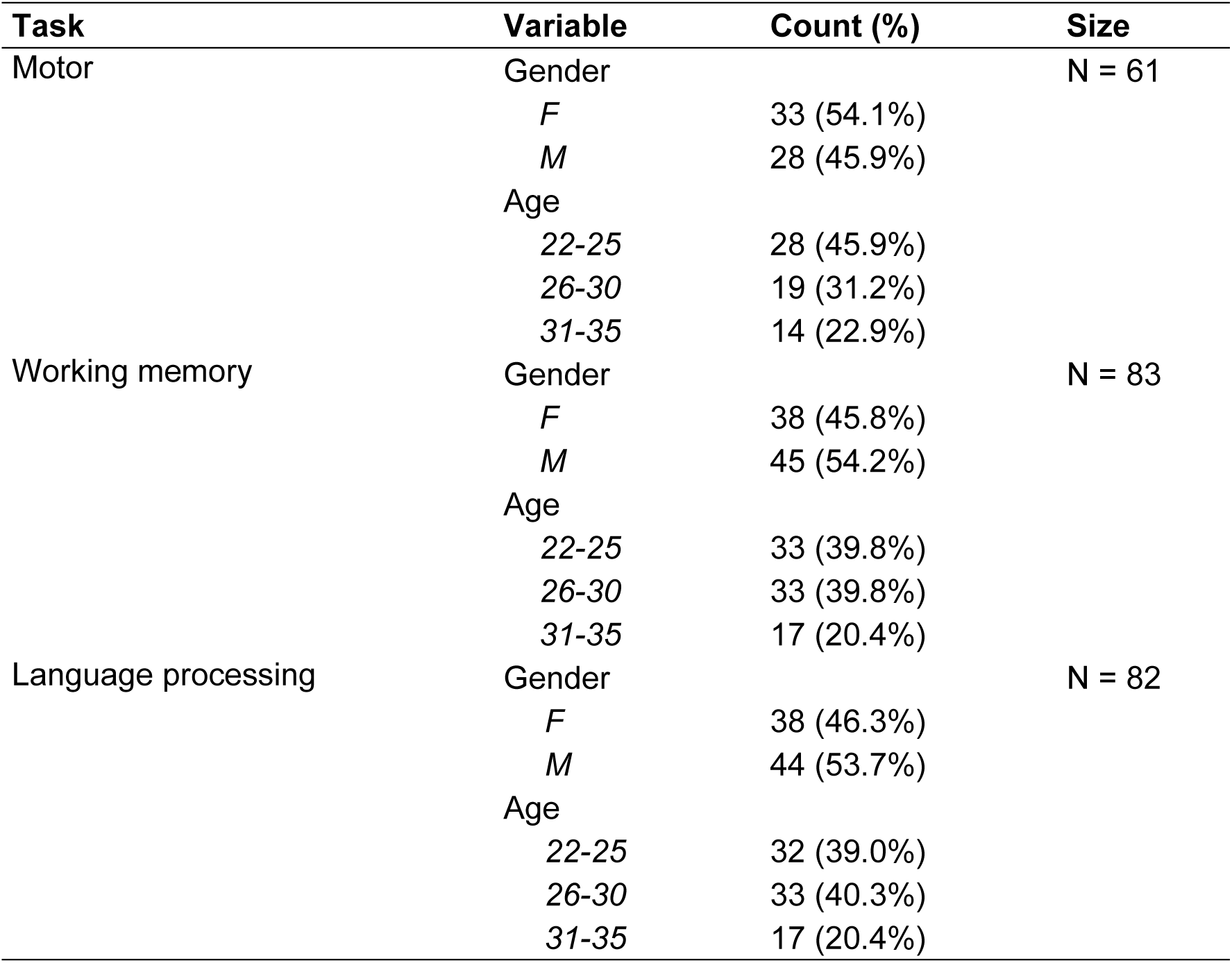
Age range and gender distribution of the study sample by task.

### 2.2. Experimental tasks

Visual and auditory stimuli for the experimental tasks were generated using E-Prime 2.0 Pro software. Visual stimuli were displayed via an LCD projector and reflected onto a mirror positioned approximately three feet above the participant’s eyes. Auditory stimuli were delivered binaurally using an MEG-compatible audio system. Participants’ responses were collected using a fiber-optic button box and recorded as a response channel within the MEG data. Further details regarding the experimental protocols can be found in the 1200 Subjects Data Release Reference Manual (WU-Minn Consortium Human Connectome Project, 2017b).

#### 2.2.1. Motor task

Sensory-motor function was evaluated using a visually cued motor task, adapted from prior protocols (Buckner et al., 2011; Thomas Yeo et al., 2011). Participants were instructed to perform repetitive movements—either tapping the thumb and index finger or squeezing the toes—based on visual cues indicating Left/Right Hand/Foot. Each movement block lasted 15 seconds and began with a 3-second instruction cue (e.g., “Right Hand”), followed by 10 paced movements. Across two runs, participants completed 32 movement blocks (8 per limb), interleaved with nine 15-second Fixation blocks (see Figure 1A).

**Figure 1.**
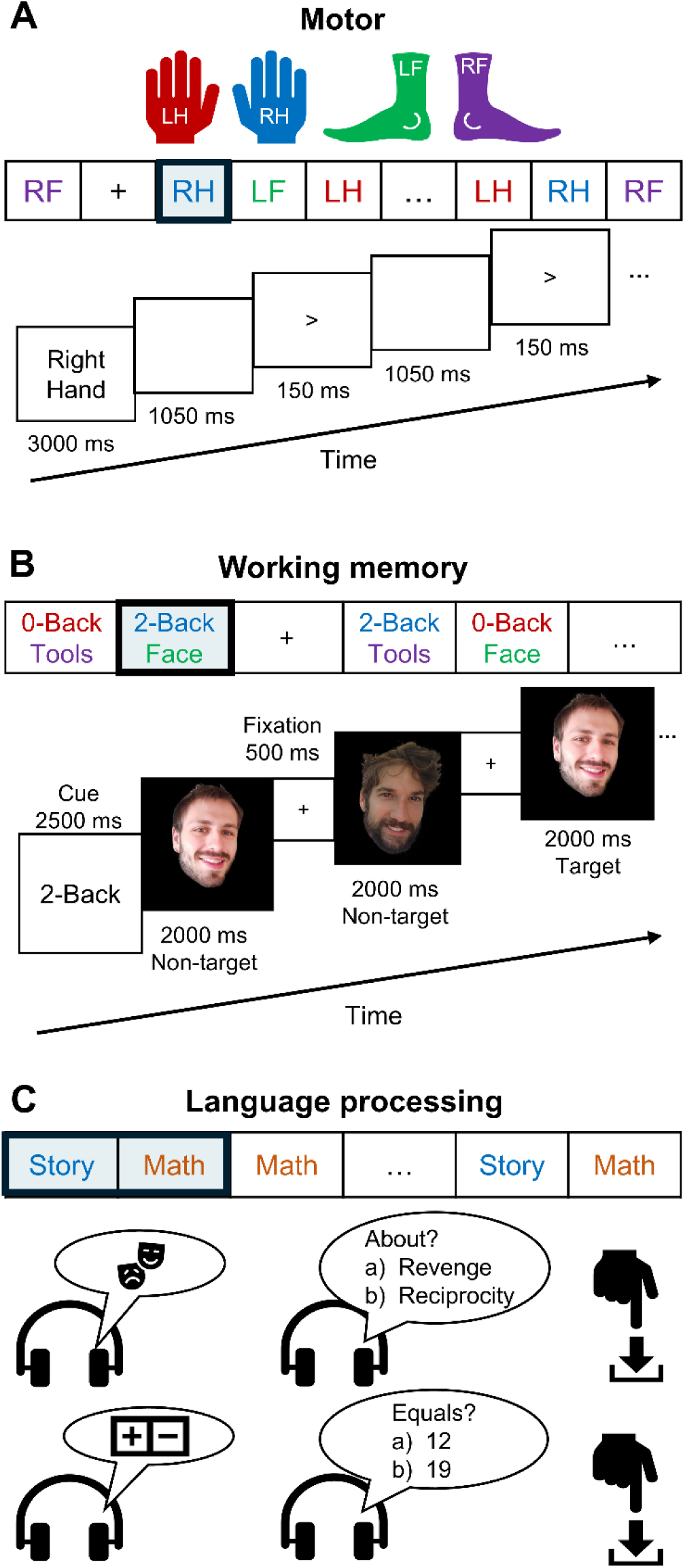
Schematic illustration of the experimental tasks. **(A)** The motor task consisted of alternating blocks of Right Hand (RH), Left Hand (LH), Right Foot (RF), and Left Foot (LF) movements, interleaved with Fixation (+) periods. An example sequence of stimuli from a Right Hand (RH) block is shown. **(B)** The working memory task comprised alternating task blocks (0-Back/2-Back and Faces/Tools) and Fixation (+) periods. The lower panel shows an example sequence of stimuli from a 2-Back/Faces block. *Note: Photographs correspond to authors J.J.H-M and E.S. and were not used as stimuli in the original HCP study.* **(C)** In the language processing task, Story and Math blocks were interleaved. In both conditions, stimuli were presented aurally, and participants provided a motor response to choose between two possible answers, corresponding either to the topic of a story or to the solution of an arithmetic problem.

#### 2.2.2. Working memory task

Working memory was evaluated using a visual N-back paradigm with sequentially presented images. Participants completed two runs of 16 blocks each, alternating between Face and Tool image blocks as well as between 0-Back and 2-Back blocks. In the 0-Back condition, participants responded when the current image matched a predefined target; in the 2-Back condition, they responded when the image matched the one presented two items earlier. Responses were made via button press using the right hand: the index finger for matches and the middle finger for non-matches. As in the motor task, task blocks were interleaved with 15-second Fixation periods (see Figure 1B).

#### 2.2.3. Language processing task

The Story-Math task was originally developed by Binder et al. (2011) and consisted of two runs alternating between blocks of narrative comprehension and mental arithmetic. Each run included 7 Story blocks and 15 Math blocks, with an average block duration of approximately 30 seconds. All stimuli were presented aurally.

In the Story condition, participants listened to short auditory narratives adapted from Aesop’s fables, followed by a two-option multiple-choice question assessing story comprehension (e.g., identifying whether a story was about *revenge* or *reciprocity*). In the Math condition, participants heard a sequence of addition and subtraction problems (e.g., “*Eight plus eleven equals*…”), followed by a choice between two possible answers (e.g., “…*twelve* or *nineteen*?”). Task difficulty in the Math condition was dynamically adjusted to maintain a consistent level of difficulty across participants. Responses were recorded via button press, with participants using the right index finger to select the first choice and the right middle finger to select the second option (see Figure 1C).

### 2.3. Data acquisition

Brain activity was recorded using a whole-head MAGNES 3600 system (4D Neuroimaging, San Diego, CA), located in a magnetically shielded room at the Saint Louis University medical campus. The system comprises 248 magnetometers and 23 reference channels. Recordings were sampled at ∼2034 Hz with a 400 Hz bandwidth, and a DC high-pass filter. MEG data for each participant were acquired in a single ∼3-hour session.

Electrooculography (EOG), electrocardiography (ECG), and electromyography (EMG) were simultaneously recorded and synchronized with the MEG signal. EOG and ECG electrodes were used to monitor ocular and cardiac activity, facilitating offline artifact removal, while EMG recordings were used to identify movement onset during the motor task.

### 2.4. Pre-processing

Pre-processed, artifact-free MEG data were obtained directly from the HCP database. The detailed pre-processing pipeline can be found in the HCP Reference Manual (WU-Minn Consortium Human Connectome Project, 2017b). Initial steps included the detection of bad channels and bad segments, followed by independent component analysis (ICA) for artifact identification and removal.

Data were segmented into 1.2-second trials time-locked to the cognitive activity of interest. In the motor task, trial onset was defined by the onset of the EMG signal recorded from hand or foot muscles. In the working memory task, trial onset corresponded to the presentation of the image that participants were required to match with the target image. In the language processing task, the first 24 seconds of each Story/Math block were segmented into consecutive 1.2-second trials.

### 2.5. Brain maps of natural frequencies

As mentioned above, this study aimed to investigate whether the brain’s natural frequencies are modulated during cognitive tasks, including motor, working memory, and language. To this end, single-subject maps of natural frequencies were generated for each participant and experimental condition and were subsequently subjected to statistical comparison (see Figure 2 for the full analysis pipeline). As described in the following sections, the analysis began with the reconstruction of source-level brain activity, followed by spectral analysis of the reconstructed signals. The resulting power spectra were then subjected to k-means clustering, allowing the identification of the most characteristic brain frequency per participant and condition on a voxel-by-voxel basis. All analyses were performed using FieldTrip version 20230118 (Oostenveld et al., 2011) and in-house Matlab code. The scripts required to reproduce the analyses and figures are available at https://github.com/necog-UAM.

**Figure 2.**
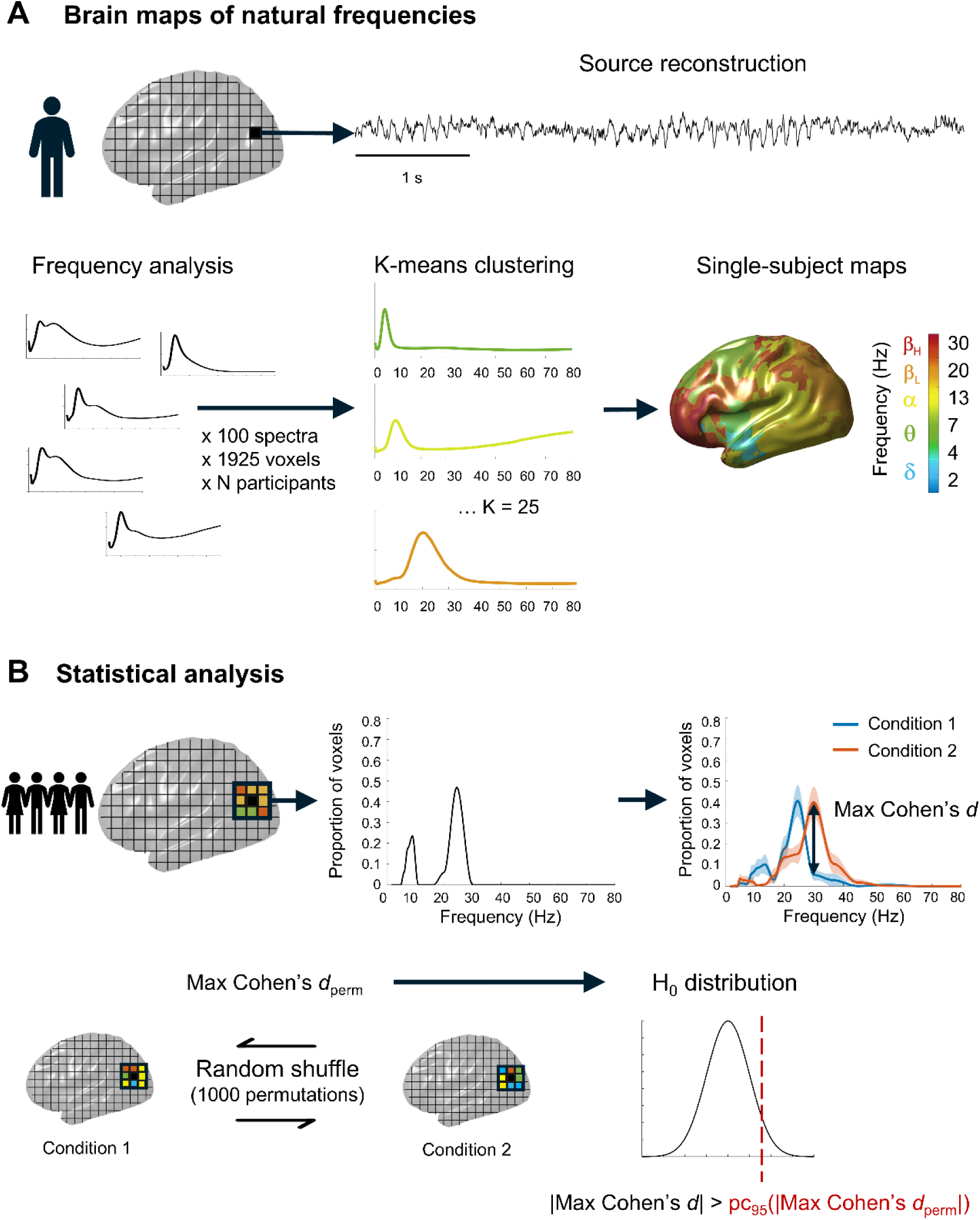
Schematic overview of the analysis pipeline. **(A)** Generation of single-subject brain maps of natural frequencies. Source-level time series were reconstructed and subsequently subjected to frequency analysis. The resulting power spectra were then classified using k-means clustering, allowing identification of the most characteristic oscillatory frequency for each participant, condition, and voxel. **(B)** Statistical analysis of brain maps of natural frequencies. For each voxel and condition, we quantified the proportion of neighboring voxels (within a 1.5 cm radius) that exhibited each oscillatory frequency. Cohen’s *d* was computed for each frequency, and the maximum *d* was retained as the test statistic. Statistical significance was assessed using permutation testing with a maximum-statistic approach to control for multiple comparisons.

#### 2.5.1. Reconstruction of source-level activity

Each participant’s T1-weighted MRI was aligned to the MEG head coordinate system using the transformation matrices provided by the HCP Data Release (WU-Minn Consortium Human Connectome Project, 2017a).

The forward model was computed using a realistic single-shell volume conductor model (Nolte, 2003). A standard MNI grid with 1 cm resolution was then warped to each participant’s individual brain anatomy, and lead fields were calculated for all voxels within the cortical and hippocampal regions (1925 voxels). Source-level time series were estimated using a linearly constrained minimum variance (LCMV) beamformer (Van Veen et al., 1997). Spatial filter weights were derived from the covariance matrix of artifact-free data, applying a regularization factor (lambda) of 10%. These beamformer weights were subsequently used to project sensor-level data into source-space time series. Because beamformer source estimates exhibit a center-of-head bias that spuriously enhances activity in deeper structures compared to cortical regions (Shapira Lots et al., 2016), we normalized each voxel’s source signal by dividing it by its standard deviation across time.

#### 2.5.2. Frequency analysis of source-level data

Spectral analysis was carried out on the reconstructed source-space data. For each condition, the epoched data were analyzed using a Hanning-tapered Fourier transform. Power estimates were obtained for 82 logarithmically spaced frequency bins spanning 1.7 to 99.5 Hz. The window length was frequency-dependent, a choice that reduces the influence of the 1/f background component and improves the detection of rhythmic activity. Given the 1.2-s trial length, the window was defined as three cycles per frequency, except for the lowest frequencies (1.7–2.6 Hz), where the number of cycles was gradually increased from 2 to 3. To ensure that all power spectra were on the same scale, each spectrum was normalized by expressing power at each frequency as a proportion of the total power across the full frequency range.

For the motor task the mean number of power spectra per voxel and participant ranged from 148 ± 7.9 (mean ± SD) to 205 ± 17.4 across conditions. For the working memory task, this range was 74 ± 7.3 to 149 ± 8.4, exceeding 100 for all conditions except Fixation. Finally, for the language processing task, the number of spectra ranged from 247 ± 11.3 to 258 ± 12.6.

#### 2.5.3. K-means clustering of power spectra

We employed a k-means clustering approach to identify distinct spectral patterns in the source-reconstructed oscillatory signals. To reduce computational demands, 100 power spectra were randomly sampled for each participant, voxel, and condition. For the Fixation condition of the working memory task, the maximum number of trials available across individuals (43 trials) was used. The selected spectra were then concatenated across all participants, brain voxels (1925), and trials (100) before clustering, separately for each task condition. The cosine distance was chosen as the similarity measure, as it emphasizes differences in spectral shape (Keitel & Gross, 2016). The number of clusters was set to 25. To ensure solution stability, the algorithm was repeated five times with a maximum of 200 iterations per run, and the solution with the lowest total within-cluster distance was retained (see Capilla et al., 2022).

#### 2.5.4. Single-subject brain maps of natural frequencies

After training a k-means clustering model for each experimental condition, individual brain maps of natural frequencies were generated following Arana et al. (2025). Although for model training, a reduced subsampling scheme (100 spectra per voxel) was maintained to limit computational cost, for the subject-level classification stage, all available spectra per voxel were utilized, thus providing more accurate estimates of individual natural frequency maps.

Thus, for every voxel and participant, the previously estimated power spectra were assigned to the group-level clusters based on the smallest cosine distance to the cluster centroids. To determine the oscillatory frequency characterizing each cluster, local peaks in the centroid power spectra were identified. Centroids lacking identifiable peaks were excluded, as spectral maxima are necessary to confirm genuine rhythmic activity (Donoghue et al., 2022). If two peaks were present, their relationship was assessed: when harmonically related (e.g., 10 and 20 Hz, ±1 Hz tolerance), only the fundamental frequency was retained; when not harmonically related (e.g., 7 and 22 Hz), each frequency was assigned an equal weight of 50%. Subsequently, the proportion of spectra assigned to each cluster was calculated and normalized using z-scores, thereby generating voxel-wise indices of the relative contribution of different oscillatory peaks (e.g., 1.9 Hz, 2.6 Hz, 3.3 Hz). To further refine the spectral representation, tenfold interpolation was applied, allowing the centroid frequencies to be expressed at a higher resolution (e.g., 1.9, 2.0, 2.1 Hz, etc.).

To improve the robustness of single-subject maps, they were spatially smoothed (see Arana et al., 2025). Conventional smoothing is not suitable in this context, as abrupt spectral transitions between neighboring voxels are common (e.g., delta and high-beta activity in frontal regions). To overcome this problem, we identified the most stable frequency across each voxel’s local neighborhood. For each voxel, neighboring voxels within a 1.5 cm radius were selected, and their z-scores representing relative oscillatory contributions were combined with those of the central voxel. At each frequency bin, one-sample t-tests against zero were performed to evaluate whether a given frequency was consistently represented across the neighborhood. The frequency yielding the highest t-value was designated as the voxel’s natural frequency, reflecting the most robust oscillatory component within the 1.5 cm radius. Voxels for which no frequency reached statistical significance (p > 0.05) were assigned missing values.

#### 2.5.5. Group-level brain maps of natural frequencies

Group-level brain maps of natural frequencies were derived from single-subject maps for visualization purposes. For each voxel, frequency distributions across participants were fitted with Gaussian functions to identify the most representative oscillatory peak. The Gaussian peak with the highest amplitude, corresponding to the frequency shared by the largest number of participants, was then selected as the voxel’s natural frequency in the group-level map.

### 2.6. Statistical analysis

Statistical differences between conditions were evaluated by performing voxel-wise comparisons of individual brain maps of natural frequencies. This type of analysis is inherently challenging because the same voxel can exhibit markedly different frequency values across participants (e.g., 5 and 25 Hz in a frontal voxel). Consequently, conventional summary statistics based on measures of central tendency, such as the mean, are inappropriate, as intermediate values (e.g., 15 Hz) would fail to capture the true variability at that voxel. To address this limitation, instead of treating the voxel’s frequency as the primary variable, we quantified the proportion of neighboring voxels (within a 1.5 cm radius) that exhibited each frequency (see Figure 2B). This approach yielded normalized local frequency distributions that describe, for each voxel, participant, and condition, the proportion of voxels in the surrounding neighborhood associated with each frequency.

Differences between experimental conditions were assessed using a non-parametric permutation testing framework combined with effect size estimation, which provides a more informative characterization of condition-related effects. Specifically, for each voxel and comparison, we computed repeated-measures Cohen’s *d* for each frequency of the normalized local frequency distributions and retained the maximum absolute *d* as the test statistic. To evaluate statistical significance, we generated a null distribution of Cohen’s *d* under the hypothesis of no condition differences by randomly shuffling condition labels. This procedure was repeated 1000 times, and in each permutation the maximum statistic was stored to control for multiple comparisons. Results were considered statistically significant if they exceeded the absolute value of the 95^th^ percentile of the null distribution.

The specific statistical contrasts conducted for each task were as follows. For the motor task, each movement condition (Left Hand, Right Hand, Left Foot, and Right Foot) was contrasted against Fixation to characterize the brain dynamics associated with movement execution. For the working memory task, each condition (0-Back, 2-Back, Faces, and Tools) was compared with Fixation to provide an overall characterization of task-related brain activity. In addition, the 0-Back and 2-Back were contrasted to isolate brain networks associated with different working memory loads, while Faces and Tools were compared to examine differences related to distinct visual and semantic processing demands. For the language processing task, the Story condition was contrasted with the Math condition to capture differential brain dynamics underlying language comprehension and mental arithmetic. For visualization purposes, we selected a subset of voxels showing statistically significant effects, focusing on local maxima in the statistical maps and their spatially symmetric counterparts.

## 3. Results

### 3.1. Motor task

Group-level cortical maps of natural frequencies for the different conditions of the motor task are shown in Figure 3A. Each brain view (left and right lateral, and left and right medial) is color-coded to represent frequencies at different bands: high-beta (β_H_, red; ∼20–30 Hz), low-beta (β_L_, yellow-orange; ∼13–20 Hz), alpha (α, green-yellow; ∼7–13 Hz), theta (θ, cyan-green; ∼4–7 Hz), and delta (δ, blue; ∼2–4 Hz). During Fixation, the brain exhibited the canonical spatial distribution of natural frequencies, with slow oscillations (delta and theta) predominating in medial fronto-temporal regions, alpha-band activity characterizing parieto-occipital cortex, and beta oscillations localized to lateral frontal areas (Capilla et al., 2022). During unilateral hand and foot movements, the overall brain pattern remained similar, but beta-band activity became more widespread across sensorimotor cortices.

**Figure 3.**
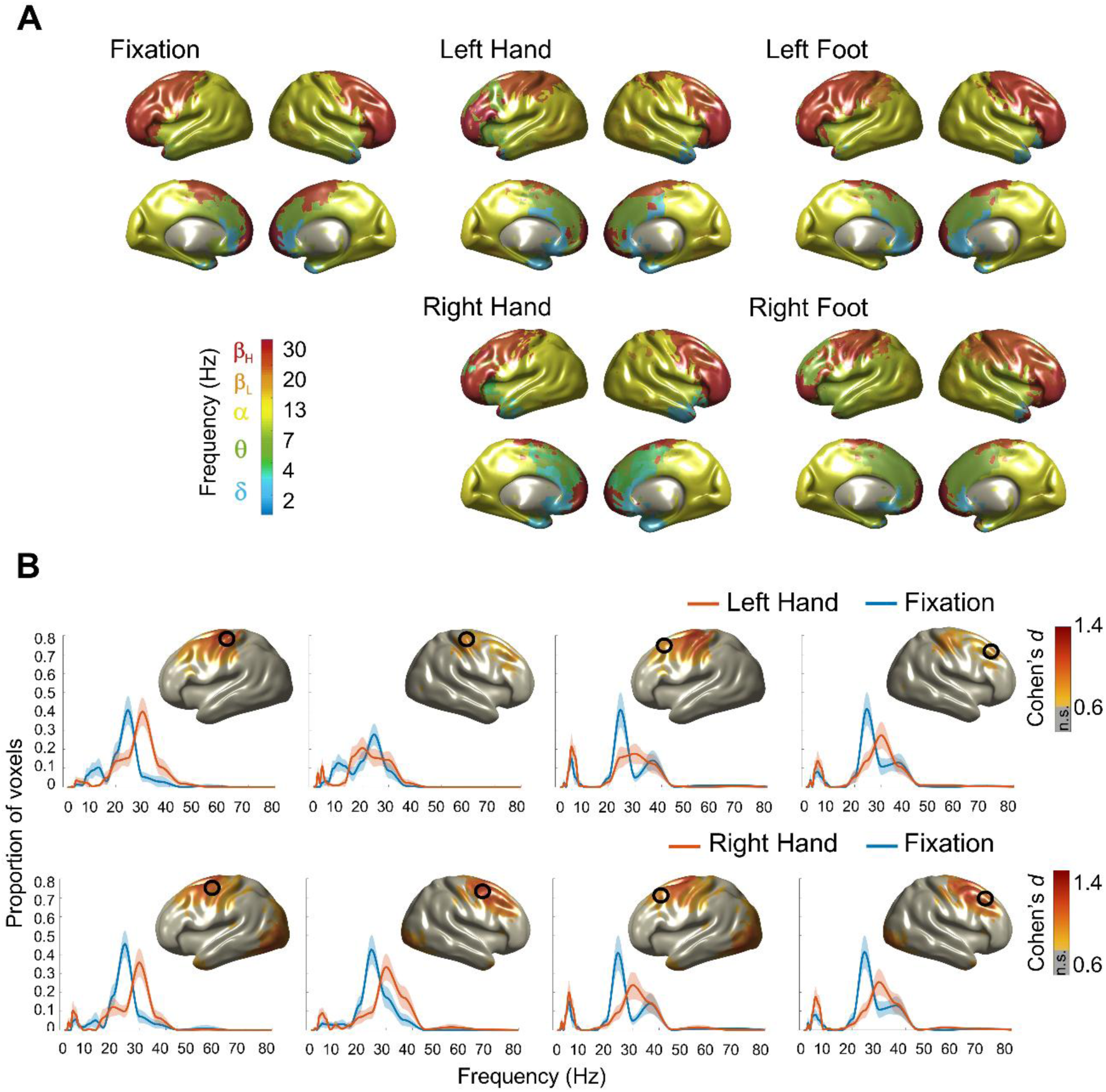
Motor task results. **(A)** Group-level cortical maps of natural frequencies for the Fixation condition and for unilateral Hand and Foot movements, illustrating the spatial distribution of characteristic oscillatory frequencies across the cortex. **(B)** Voxel-wise contrasts between Left/Right Hand and Fixation, showing the cortical regions where natural frequencies differed between conditions. Frequency distributions were extracted from selected representative voxels to illustrate condition-specific shifts in oscillatory activity. Mean ± Standard Error of the Mean (S.E.M.) across subjects is shown for each condition. Effect size cortical maps corresponding to the contrasts depict the magnitude of spectral differences across cortical regions (non-significant results are displayed in grey, n.s.).

In general, the statistical contrasts between Left/Right Hand and Fixation revealed a significant acceleration in the typical beta-band activity in bilateral sensorimotor and premotor cortices (Figure 3B). While beta-band oscillations peaked at approximately 24 Hz during Fixation, this increased to ∼30 Hz during movement execution. In the Left Hand condition, this rightward shift was accompanied by a concomitant disappearance of alpha-band oscillations in bilateral sensorimotor cortices. The magnitude of the effect exceeded Cohen’s *d* > 1 in sensorimotor regions, indicating large effect sizes. Notably, the effects were symmetrical and consistent across hand conditions. Residual delta/theta-band activity was also observed in motor and premotor cortices in both hand conditions, although the differences relative to Fixation did not reach significance.

In contrast to the robust hand-related pattern, comparisons between Left and Right Foot and Fixation revealed very small, statistically significant effects confined to isolated voxels.

### 3.2. Working memory task

During the Fixation periods of the working memory task, the group-level map also exhibited the characteristic pattern of natural frequencies at rest, similar to the Fixation condition of the motor task (see Figure 4A). In contrast, the task-related condition maps showed an overall shift toward lower frequencies (from alpha/beta to delta/theta) over left motor regions.

**Figure 4.**
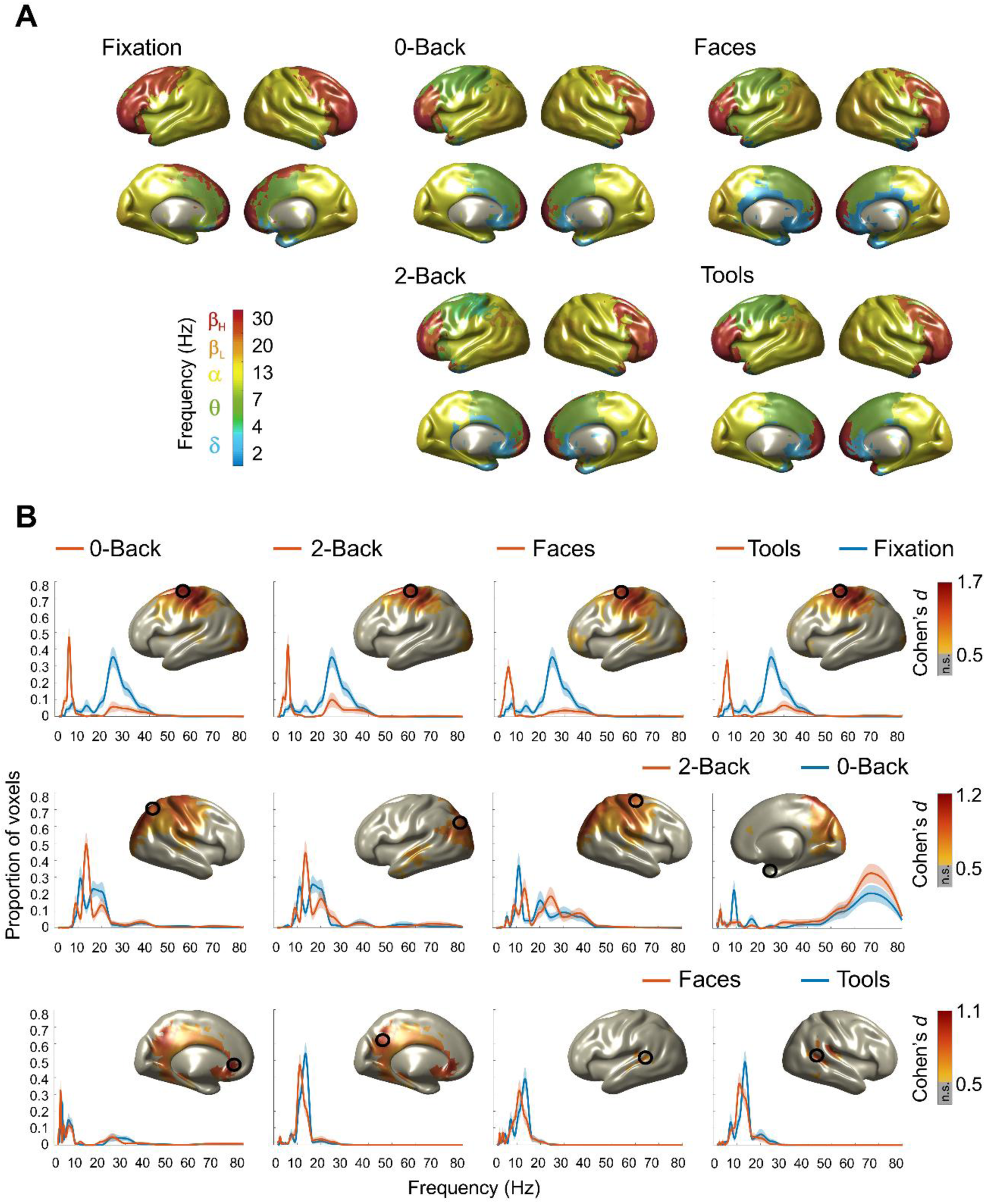
Working memory task results. **(A)** Group-level cortical maps of natural frequencies across the different conditions of the working memory task, showing the spatial distribution of characteristic oscillatory frequencies throughout the cortex. **(B)** Voxel-wise contrasts illustrating: (top) differences between each experimental condition and Fixation, (middle) the contrast between 2-Back and 0-Back conditions to isolate memory load effects, (bottom) the contrast between Faces and Tools conditions to isolate item-specific effects. Frequency distributions were extracted from selected representative voxels to illustrate condition-specific shifts in oscillatory activity. Mean ± S.E.M. across subjects is shown for each condition. Effect size cortical maps corresponding to the contrasts depict the magnitude of spectral differences across cortical regions (non-significant results are displayed in grey, n.s.).

Comparisons of each task-related condition with Fixation revealed a very large and noticeable effect (Cohen’s *d* > 1.5) in motor regions of the left hemisphere (Figure 4B). In all cases, oscillatory activity in the left motor cortex shifted from beta-band activity peaking at ∼24 Hz to theta activity peaking around 6 Hz.

To identify load-dependent changes in oscillatory activity during working memory, we contrasted the 2-Back and 0-Back conditions (Figure 4B). The most prominent effect was a shift in a bimodal alpha/beta peak in parieto-occipital areas, from approximately 11/16 Hz in the 0-Back condition to 13/20 Hz in the 2-Back condition, as well as in central regions, from 11/20 Hz to 13/24 Hz. In addition, the 2-Back condition was associated with an increase in gamma-band activity (60–80 Hz) in the medial temporal lobe, accompanied by a suppression of lower-frequency activity (<20 Hz). Overall, these spectral shifts indicate a transition from slower oscillatory dynamics during low-load maintenance (0-Back) to faster rhythmic activity during active updating (2-Back), with effect sizes that can be considered large (Cohen’s *d* > 1).

In contrast, only a limited number of significant differences were observed between Faces and Tools (Figure 4B). The most prominent effect was an acceleration of alpha-band activity, with a higher peak frequency in response to Tools (13 Hz) compared to Faces (11 Hz) in medial temporal and parietal regions.

### 3.3. Language processing task

At the group level, both the Story and Math conditions of the language processing task exhibited widespread delta-band activity across temporal and inferior frontal cortices, deviating from the canonical resting-state pattern (Figure 5A).

**Figure 5.**
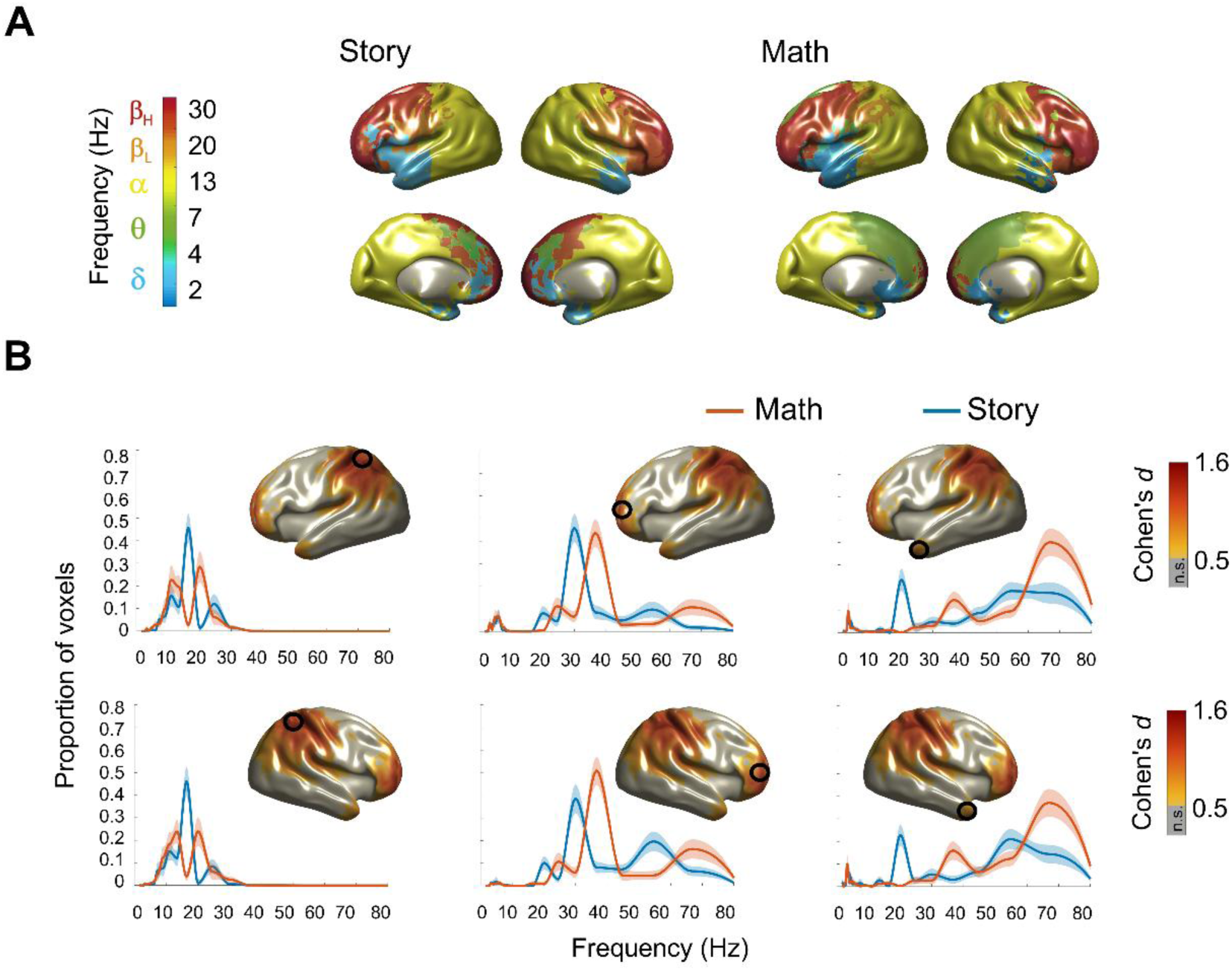
Language processing task results. **(A)** Group-level cortical maps of natural frequencies for the Story and Math conditions, illustrating the spatial distribution of characteristic oscillatory frequencies across the cortex. **(B)** Voxel-wise contrasts between Story and Math conditions, highlighting cortical regions where natural frequencies differed between the two tasks. Frequency distributions were extracted from selected representative voxels to illustrate condition-specific shifts in oscillatory activity. Mean ± S.E.M. across subjects is shown for each condition. Effect size cortical maps corresponding to the contrasts depict the magnitude of spectral differences across cortical regions (non-significant results are displayed in grey, n.s.).

The statistical contrast between Story and Matth revealed a task-dependent divergence in frequency profiles within the frontoparietal network (Figure 5B). During Math processing, high-beta and gamma peaks exhibited a rightward shift of approximately 5-15 Hz (from 20/30 Hz to 36 Hz, and from 55 Hz to ∼70 Hz), particularly in bilateral medial temporal and prefrontal regions (Cohen’s *d* > 1.5). A similar pattern was observed in parietal regions, with a ∼5 Hz acceleration (from 16 Hz to 20 Hz) in the beta band in the Math condition relative to Story.

## 4. Discussion

This study investigated whether natural frequencies constitute stable intrinsic properties of the brain or dynamically reconfigure during cognitive processing. Applying a data-driven spectral clustering approach to source-reconstructed MEG data from the HCP, we mapped natural frequencies across motor execution, working memory, and language processing tasks. Our results demonstrate that while the global spatial organization of natural frequencies remains largely preserved during task engagement, specific cortical regions exhibit systematic, task-dependent shifts in their characteristic spectral peaks. During hand movements, sensorimotor and premotor cortices shifted from low-beta (∼24 Hz) to high-beta rhythms (∼30 Hz). Working memory manipulation (2-Back vs. 0-Back) accelerated parieto-occipital alpha/beta activity and recruited high-gamma oscillations (∼60–80 Hz) in medial temporal regions. Arithmetic processing, in turn, increased beta/gamma frequencies across frontoparietal networks relative to semantic comprehension. Collectively, these findings reveal that natural frequencies reflect a hybrid architecture: globally stable yet locally flexible in response to cognitive demands.

### 4.1. Motor execution: Beta modulation beyond power

Our observation of beta-band modulations during motor execution both converges with and extends classical accounts of movement-related desynchronization and synchronization (Kilavik et al., 2013). Previous studies consistently report decreases in beta power over sensorimotor regions preceding and during movement, followed by a rebound in beta activity approximately 0.3–1 s after movement termination (Neuper & Pfurtscheller, 2001; Pfurtscheller & Lopes da Silva, 1999). By tracing shifts in characteristic frequency rather than relying solely on spectral power, the present results reveal an upward shift in oscillatory activity within the beta range itself, from ∼24 Hz during fixation to ∼30 Hz during motor execution. Given that our analysis window extends to 1.2 s after movement onset and spans the full movement cycle from execution to termination, the ∼30 Hz activity likely reflects the post-movement beta rebound rather than oscillatory dynamics strictly tied to movement execution. This interpretation is consistent with previous findings showing that beta oscillations often exhibit a slightly higher and more variable peak frequency during the post-movement beta rebound compared with baseline levels (Espenhahn et al., 2017).

The post-movement period has been proposed to support the recalibration or resetting of the motor system, allowing the sensorimotor cortex to adapt to updated sensory and motor conditions in preparation for subsequent actions (Gaetz & Cheyne, 2006). Within this framework, our findings suggest that such recalibration may involve a shift in the temporal dynamics of sensorimotor rhythms toward faster beta modes. This interpretation aligns with the emerging evidence that beta peak frequency and sub-band structure are behaviourally meaningful and may differ across motor circuits and effectors (Borra et al., 2023; Espenhahn et al., 2017). Consistent with this view, recent intracranial and non-invasive studies indicate that different beta components may support distinct aspects of motor preparation, execution, and adaptation (Jahani et al., 2020; Unterweger et al., 2020). Specifically, while low-beta (∼15–20 Hz) has often been associated with maintenance of the status quo, higher beta frequencies (∼25–35 Hz), bordering the low-gamma range, have been linked to active muscle contraction and sensorimotor integration (Echeverria-Altuna et al., 2022; Spitzer & Haegens, 2017).

### 4.2. Working memory: Effects of motor response, load, and stimulus type

#### 4.2.1. Motor-related beta suppression

In the working memory task, we observed a very robust and large effect over sensorimotor regions of the left hemisphere (Cohen’s *d* > 1.5) when comparing all working memory conditions (0-Back, 2-Back, Faces, Tools) with the fixation baseline. The natural frequency of this region exhibited a pronounced shift from the beta band (∼24 Hz) during fixation to the theta band (6 Hz) during task performance. A common feature across all N-back conditions was that participants responded using their right hand, pressing the index finger for matches and the middle finger for non-matches. Accordingly, the modulation observed over left sensorimotor regions most likely reflects the button-press movement associated with the motor response required by the task. Unlike in the motor task, the analysis window in the working memory paradigm was time-locked to stimulus onset and therefore encompassed motor preparation and execution rather than the post-movement period. Consequently, the observed effect appears to correspond to beta desynchronization, which has commonly been interpreted as indexing cortical disinhibition of task-relevant neuronal populations and a transition from a stable to a more labile motor state (Borra et al., 2023; Unterweger et al., 2020).

In addition to the suppression of beta-band oscillations over the contralateral sensorimotor cortex, we found a concomitant increase in theta-band activity. These findings are in line with recent reports of slow oscillations (∼3–5 Hz) emerging immediately before and at movement onset, coinciding with the decrease in beta power. Unlike beta desynchronization, this slow activity is transient, typically more focal and strongly lateralized, and is confined to movement onset, suggesting that it may serve as a precise temporal marker of action initiation (Bönstrup et al., 2025). During the motor task, we also detected some delta/theta activity; however, these effects were relatively weak and likely reflect residual activity, as such slow oscillations typically occur immediately before or around, rather than after, movement onset.

#### 4.2.2. Load-dependent frequency acceleration

Our results indicate that increasing working memory load was associated with a shift toward higher oscillatory frequencies in the alpha and beta ranges over parieto-occipital regions, together with the recruitment of gamma oscillations in the medial temporal lobe. In contrast, theta-band oscillations—characteristic of medial frontal regions—did not exhibit a clear load-dependent frequency effect, despite they are consistently engaged in working memory tasks (Pavlov & Kotchoubey, 2022).

Midfrontal theta power has been shown to rise with increasing memory load, reflecting greater executive control demands (Brookes et al., 2011; Jensen & Tesche, 2002; Maurer et al., 2015). Interestingly, recent evidence suggests that higher memory load may also be associated with a slight slowing of the theta peak frequency (Ratcliffe et al., 2022). This finding might provide support to the Jensen–Lisman model (Jensen & Lisman, 1998; Lisman & Jensen, 2013), which proposes that slower theta cycles may allow a larger number of gamma-nested representations to be maintained simultaneously. Although our analysis explicitly focused on frequency rather than power modulations, we did not observe the theta slowing reported by Ratcliffe et al. (2022). As noted by the authors, however, this effect is relatively small (e.g., 5.85 Hz in the 1-back vs. 5.77 Hz in the 2-back condition in their study), and our approach may have lacked the sensitivity required to reliably detect such subtle frequency shift.

In contrast to the absence of theta effects, we found an acceleration of approximately 2–4 Hz in the natural frequencies of parieto-occipital cortex within the alpha and beta ranges as working memory demands increased. Previous studies have reported mixed effects of alpha and beta power in relation to memory load, with some showing increases and others decreases, particularly in visual working memory tasks like the one used here (Pavlov et al., 2022). In line with our findings, however, prior work has reported higher alpha peak frequencies in posterior regions under greater memory demands. This shift toward faster alpha rhythms has been interpreted as reflecting accelerated cycles of cortical inhibition and enhanced temporal resolution of neural processing, thus enabling cortical networks to sustain and coordinate a larger number of active representations (Haegens et al., 2014; Sghirripa et al., 2021).

Finally, high-gamma activity (60–80 Hz) in medial temporal lobe regions was significantly more consistent in the 2-Back relative to the 0-Back condition. This recruitment of gamma during active manipulation aligns with recent evidence linking high-frequency oscillations to item-specific maintenance and hippocampal-cortical communication during working memory tasks (Carver et al., 2019; Costers et al., 2020; Mirjalili et al., 2025). Importantly, our data-driven approach revealed this effect without a priori band selection, thereby reducing the risk of misattributing broadband aperiodic activity to rhythmic gamma oscillations (Donoghue et al., 2022).

#### 4.2.3. Subtle stimulus category effects

Regarding stimulus specificity, we observed only subtle differences between Faces and Tools conditions, primarily limited to a slight alpha acceleration for Tools. While fMRI studies robustly identify category-specific regions like the fusiform face area (Barch et al., 2013; Kanwisher et al., 1997), our spectral analysis suggests that oscillatory frequency may be less sensitive to stimulus category than to process-related factors (e.g., memory load), or that category-specific spectral fingerprints are more subtle than load-dependent reconfigurations. These findings are consistent with recent MEG work indicating that category information is often encoded in spatial patterns of activity rather than in gross spectral shifts (Kaiser et al., 2016).

### 4.3. Language processing: Semantic integration vs. arithmetic computation

The language processing task revealed that both Story and Math conditions elicited widespread delta-band activity over temporal and inferior frontal cortices, diverging from characteristic resting-state or fixation patterns (Capilla et al., 2022). This delta predominance aligns with evidence that slow oscillations (<4 Hz) track higher-order linguistic structure, phrasal integration, and narrative-level processing (Lu et al., 2023; Molinaro & Lizarazu, 2018; Rimmele et al., 2021; Slaats et al., 2023; Ten Oever et al., 2022). Furthermore, delta-band tracking in temporal and inferior frontal regions has been linked to syntactic grouping, predictive coding of multi-word chunks, and top-down constraints from frontal to auditory cortex during speech comprehension (Lu et al., 2023; Mai & Wang, 2023; Rimmele et al., 2021; Slaats et al., 2023; Ten Oever et al., 2022). This suggests that both Story and Math tasks, involving auditory-verbal processing, recruited core language–comprehension networks beyond a canonical resting-state configuration, supporting semantic integration of auditory information.

In contrast, arithmetic reasoning recruited higher-frequency beta/gamma activity across frontoparietal networks compared to narrative comprehension. This spectral acceleration is consistent with evidence that demanding numerical operations rely on frontoparietal beta–gamma interactions, including strengthened beta-band connectivity in the right parietal cortex during magnitude comparison (González-Garrido et al., 2018) and theta–gamma or high-gamma upregulation during working memory tasks (Carver et al., 2019).

Moreover, it has been reported that numerical manipulation elicits alpha/beta-band (8-18 Hz) power reduction in bilateral superior parietal cortices (Koshy et al., 2020), which aligns with our 16 Hz suppression and 20 Hz increment in the Math condition. In a similar vein, the increased gamma-band activity observed in the Math condition is in line with recent findings linking parietal gamma oscillations (30–40 Hz) to the processing of symbolic arithmetic proofs and mathematical expertise (Gashaj et al., 2024).

These results extend prior frameworks indicating that beta power increments might reflect the active maintenance of a neurocognitive network configuration, whereas beta power decreases may indicate network revision, as observed during semantic unification (Bastiaansen & Hagoort, 2006; Lewis & Bastiaansen, 2015). Thus, while the beta/gamma acceleration during arithmetic may reflect the rapid, sequential updating required for mental calculation, the slower beta/gamma interaction during semantic processing may reflect the active maintenance and integration of information. The co-occurring suppression of lower frequencies (<30 Hz) in temporal regions further suggests a trade-off between slower integrative rhythms and faster, transient processing modes during executive control—a subtle effect obscured when analyses rely on predefined frequency bands rather than frequency-resolved approaches. Notably, these effects emerged from direct frequency comparisons rather than power-based metrics, offering a complementary perspective on how spectral dynamics support distinct cognitive operations.

### 4.4. Theoretical implications: Frequency acceleration as a marker of cognitive engagement

A fundamental contribution of this work is the characterization of a general principle: cognitive engagement tends to accelerate neural rhythms. While the global spatial gradient of natural frequencies (slow posterior/medial to fast anterior/lateral) remains stable, local nodes shift toward higher frequencies as task demands increase. This observation supports a growing body of evidence linking oscillatory speed to arousal, attention, and processing demands (e.g. Rodriguez-Larios et al., 2020; Rodriguez-Larios & Alaerts, 2019).

This framework suggests that slow bands (delta, theta) are associated with lower arousal or integrative states (e.g., deep sleep, relaxation, semantic integration), while higher frequencies (beta, gamma) relate to top-down control, attention, and active processing (Anand et al., 2024; Rodriguez-Larios et al., 2020; Rodriguez-Larios & Alaerts, 2019). For instance, during attentional tasks, alpha power decreases in task-relevant regions (reflecting release from cortical inhibition) while gamma power increases (Fries et al., 2001; Womelsdorf et al., 2006). Our results provide macroscopic evidence of this principle in humans: the brain effectively “speeds up” its operational frequency in regions specialized for the task at hand. This has important implications for the communication-through-coherence theory as shifts in frequency alters the temporal window for neuronal communication and may affect the information coding capacity of the network (Fries, 2015).

Recent literature on neural timescales supports this view, proposing that cortical regions operate on intrinsic timescales that are functionally dynamic. Thus, neuronal timescales are shaped by cortical microarchitecture and can expand or contract based on task demands (Gao et al., 2020). Similarly, Hasson et al. (2015) proposed a hierarchy of temporal receptive windows, where sensory areas process fast-changing inputs while associative regions integrate information over longer periods. Our findings extend this framework by showing that even within these established hierarchies, local frequencies can shift upward to accommodate immediate processing requirements. This aligns with recent proposals that neuronal sequences contract or dilate as behavior speeds up or attention increases, mediated in part by neuromodulators such as acetylcholine (Buzsáki, 2026). Inhibitory interneurons, the main targets of acetylcholine, adjust their firing frequency with changes in speed and attentional demand, suggesting a plausible subcortical mechanism underlying the spectral accelerations we observed at the cortical level.

### 4.5. Conclusion

In conclusion, we sought to determine whether natural frequencies constitute fixed cortical properties or flexible dynamics shaped by cognitive context. Our results support a hybrid solution: the brain’s spectral architecture exhibits a stable global scaffold—preserving the posterior-to-anterior and medial-to-lateral gradients observed at rest—while allowing local, task-appropriate tuning of natural frequencies within functionally relevant networks. Moreover, our findings point to a general principle whereby cognitive engagement tends to accelerate neural rhythms.

By moving beyond band-limited power analyses to track frequency reconfiguration directly, we provide evidence that the cortex preserves its spectral identity while flexibly adjusting its temporal operating mode—a principle that may generalize across diverse cognitive domains. Future work leveraging this framework could illuminate how spectral reconfiguration breaks down in neurological and psychiatric conditions, potentially revealing novel biomarkers rooted not in *how much* power changes, but in *how fast* the brain oscillates when it matters most, thereby contributing to a more refined understanding of the spectral architecture of human brain function.

## Acknowledgements

We are thankful to the researchers and technicians involved in The Human Connectome Project (HCP) for recording and making publicly available the database employed in the present study. The authors also thank Jorge San Segundo for valuable advice regarding the manuscript.

This work was supported by Ministerio de Ciencia e Innovación / Agencia Estatal de Investigación, Spain / FEDER/FSE+, UE (MCIN/AEI/10.13039/501100011033/FEDER/FSE+, UE; grants PRE2022-101613 to ES, and PID2021-125841NB-I00 and PID2024-161032NB-I00 to AC).; and the Comunidad de Madrid, Spain (IND2022/SOC-23652 to LA and PIPF-2024/SAL-GL-34549 to J.J.H-M).

## Author contributions

J.J.H-M.: Conceptualization, Data curation, Formal analysis, Methodology, Software, Visualization, Writing – original draft preparation, Writing – review & editing; E.S.: Formal analysis, Methodology, Writing – review & editing; L.A.: Methodology, Resources, Writing – review & editing; A.C.: Conceptualization, Data curation, Funding acquisition, Methodology, Writing – original draft preparation, Writing – review & editing.

